# Excess Mortality Related to Chikungunya Epidemics in the Context of Co-circulation of Other Arboviruses in Brazil

**DOI:** 10.1101/140491

**Authors:** André R. R. Freitas, Luciano P. G. Cavalcanti, Andrea P. B. von Zuben, M. Rita Donalisio

## Abstract

Chikungunya is an emerging arbovirus that reached the Western Hemisphere at the end of 2013. Studies conducted in Indie Ocean and India suggest that many of deaths associated with chikungunya cannot be recognized by passive surveillance system. The occurrence of these deaths can be inferred by an increase in the overall mortality observed during chikungunya epidemics. To evaluate the mortality associated with chikungunya epidemics in Brazil, we studied monthly mortality by age group comparing the pre-chikungunya period with the chikungunya epidemic period in the most affected states of Brazil. We obtained official data from National System of Notifiable Diseases (SINAN) and Mortality Information System (SIM), both maintained by the Ministry of Health. It was possible to identify a significant increase in all-cause mortality rate during chikungunya epidemics, there was no similar mortality in previous years, even during dengue epidemics. We estimated an excess of 4,842 deaths in Pernambuco during the chikungunya epidemics (51.4/100,000 inhabitants), the most affected age groups were the elderly and under 1 year of age, the same pattern occurred in all states. Further studies at other sites are needed to confirm the association between increased mortality and chikungunya epidemics. If these findings are confirmed, it will be necessary to revise guidelines to recognize the real mortality associated with chikungunya and to improve therapeutic approaches and protective measures in the most vulnerable groups.

## Introduction

On December 9, 2013, the Pan American Health Organization (PAHO) has issued a alert about the transmission of chikungunya in the Americas(1). Since then, the transmission of this virus has been confirmed in 44 countries and territories in the region, with more than 2 million reported cases and 403 deaths. In Brazil the transmission was confirmed in September 2014, but the worst epidemic occurred in 2016 with 216,102 cases reported in 696 municipalities and 91 deaths. The lethality observed in Brazil and the rest of the Americas, has been much lower than that reported in other countries (1), perhaps due to difficulty in diagnosis and lack of adequate tools for surveillance (1). The objective of this study was to evaluate the mortality associated with chikungunya epidemics in Brazil by comparing the mortality of pre-chikungunya periods with mortality during chikungunya epidemics (2,3).

## Methods

To evaluate the possible effect of chikungunya on all-cause mortality, we compared the number of deaths expected for the current estimated population with that observed during the epidemic. The expected mortality rates were calculated based on the mean monthly Age Standardized Mortality Rate (ASMR, using the Brazilian population, census 2010) of the same months between 2011 to 2013, years prior to chikungunya circulation. Values conservative 99% of the upper limit of the confidence interval were calculated for each month as a maximum threshold of the monthly-expected deaths for each studied region. We defined excess deaths as the difference between the observed and expected deaths during the months when the mortality was higher than the threshold.

We also compared the expected and observed ASMR for each month during the chikungunya epidemic period and calculated change in mortality rate per 100,000 and percentage change (%) in each region. Similar methodology has been used previously(2–4). In most affected areas we also calculated the monthly mortality rate and 99% of the upper limit of the confidence interval by age group. As expected mortality rates we used the mean monthly mortality rate by age group of the same months of the non-epidemic years 2011-2013. Observed mortality rates were calculated for each month from January 2015 to April 2016. We studied the 3 Brazilian states with higher chikungunya incidence rate(IR) Pernambuco(9,410,772 inhabitants), Rio Grande do Norte (3,474,998 inhabitants), and Bahia(15,498,733 inhabitants) (5). Bahia is the largest and most populated state, with a heterogeneous incidence of chikungunya, and therefore we analyzed the 9 administrative regions of Bahia.

We obtained population data (census and projections) from the official Brazilian Institute of Geography and Statistics (IBGE). The mortality data were obtained from the official Mortality Information System. Cases of probable chikungunya, dengue, and zika virus (laboratory and clinical epidemiological criteria) were obtained from the Brazilian Epidemiological Surveillance System(SINAN), based on cases reported by healthcare professionals. For the processing of data and figures and the statistical analysis, we used Excel-2013^®^ and SPSS v. 20.

## Results

During the period of the chikungunya epidemic (Table 1) there was an increase in allcause mortality in all analyzed states (Figure 1). In all regions, the excess deaths occurred almost simultaneously with the peak of the chikungunya epidemic (Figure 1). The mortality were higher exactly where were the highest the chikungunya incidence rate too, we found a positive correlation between change in mortality rate and IR of CHIKV for each area (Pearson Correlation R = 0.66, p<0.03). Although cases of dengue and zika virus infection emerged concomitantly in the studied region, there were no statistically significant correlations between change in mortality rates and IR associated to these arboviruses (p = 0.14, p = 0.46, respectively). Incidence of dengue virus was high in all studied areas in the years 2011-2013 used as a baseline (5). In the state of Pernambuco, the highest incidence of dengue occurred in 2015, a year in which there were no excess deaths (Figure 1). Therefore we suggest that the excess deaths should be related to chikungunya infections and not another arbovirus that occurs at the same season.

**Figure 1.**
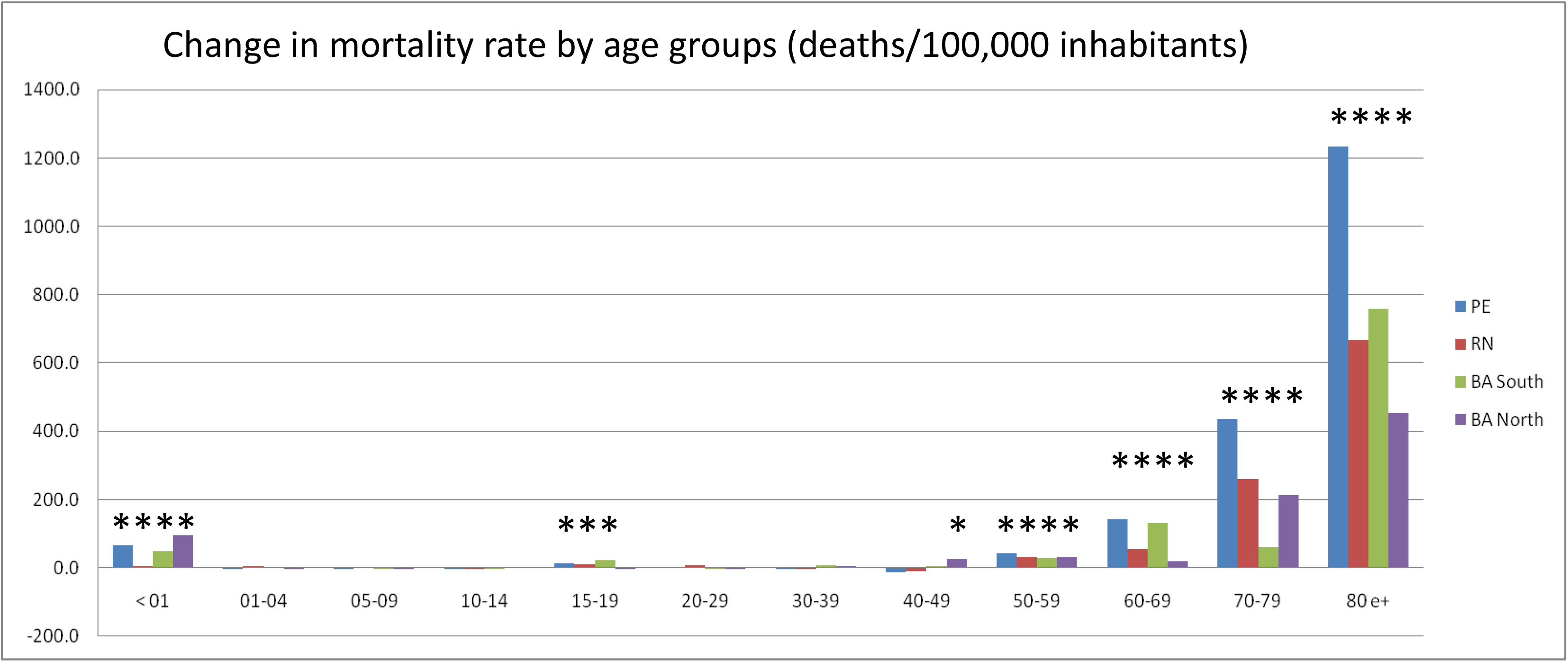
Observed and expected deaths with 99% confidence interval and incidence rate of chikungunya and dengue in Pernambuco, Rio Grande do Norte and Bahia (two regions).

**Table 1.** Observed and expected deaths, all-causes mortality rates (deaths/100.000 inhabitants), excess mortality, change in mortality rate and incidence rate of chikungunya, zika, and dengue virus in 3 states of Northeast Brazil.

|  | Population 2016 | Epidemic period | Observe deaths, (epidemic period) | Expected death, (epidemic period) | Excess death for epidemic period | Change in mortality rate for epidemic period (per 100,000) | % Change in mortality rate | CHIKV during epidemic period (cases/100,000) | ZIKA during epidemic period (cases/100,000) | DENV during epidemic period (cases/100,000) |
| --- | --- | --- | --- | --- | --- | --- | --- | --- | --- | --- |
| Pernambuco | 9,410,772 | nov/15-apr/16 | 33,677 | 28,836 | <u>4842</u> | 51.4 | <u>17%</u> | 422.1 | 3.9 | 702.1 |
| Rio Grande do Norte | 3,474,998 | feb-apr/16 | 5,378 | 4,559 | <u>819</u> | 23.6 | <u>18%</u> | 481.1 | 81.6 | 1646.4 |
| Bahia |  |  |  |  |  |  |  |  |  |  |
| East Center | 2,316,941 | feb-mar/16 | 2,076 | 1,945 |  | 5.6 | 7% | 356.8 | 195.6 | 762.4 |
| North center | 843,759 | feb-mar/16 | 721 | 672 | <u>48</u> | 5.7 | <u>7%</u> | 102.9 | 427.1 | 693.9 |
| Far South | 859,824 | feb-apr/16 | 845 | 844 |  | 0.1 | 0% | 46.9 | 270.4 | 593.6 |
| East | 4,910,439 | feb-mar/16 | 4,400 | 4,794 |  | -8.0 | -8% | 20.1 | 50.8 | 402.6 |
| Northeast | 899,179 | jan-mar/16 | 1,285 | 1,118 | <u>167</u> | 18.6 | <u>15%</u> | 210.7 | 58.2 | 349.2 |
| North | 1,127,789 | feb-mar/16 | 1,117 | 901 | <u>215</u> | 19.1 | <u>24%</u> | 457.9 | 226.5 | 910.0 |
| West | 989,288 | mar-apr/16 | 791 | 753 |  | 3.9 | 5% | 9.4 | 30.4 | 551.8 |
| South-west | 1,844,304 | feb-mar/16 | 1,596 | 1,473 | <u>123</u> | 6.7 | <u>8%</u> | 26.2 | 240.0 | 630.3 |
| South | 1,707,209 | fev-apr/16 | 3,013 | 2,543 | <u>470</u> | 27.5 | <u>18%</u> | 1178.1 | 1232.4 | 3320.9 |

Locations where the number of deaths was higher than the 99% confidence interval at least a month

The change in mortality rates(%) and the corresponding excess deaths in these 4 critical regions were: Pernambuco (16.8% and excess death = 4,842), Rio Grande do Norte (18.0% and 819), South Bahia (21.1% and 470), and North Bahia (23.9% and 215) Table 1). Figure 2 and Table 2 show excess mortality rates by age group for each region.

**Figure 2.**
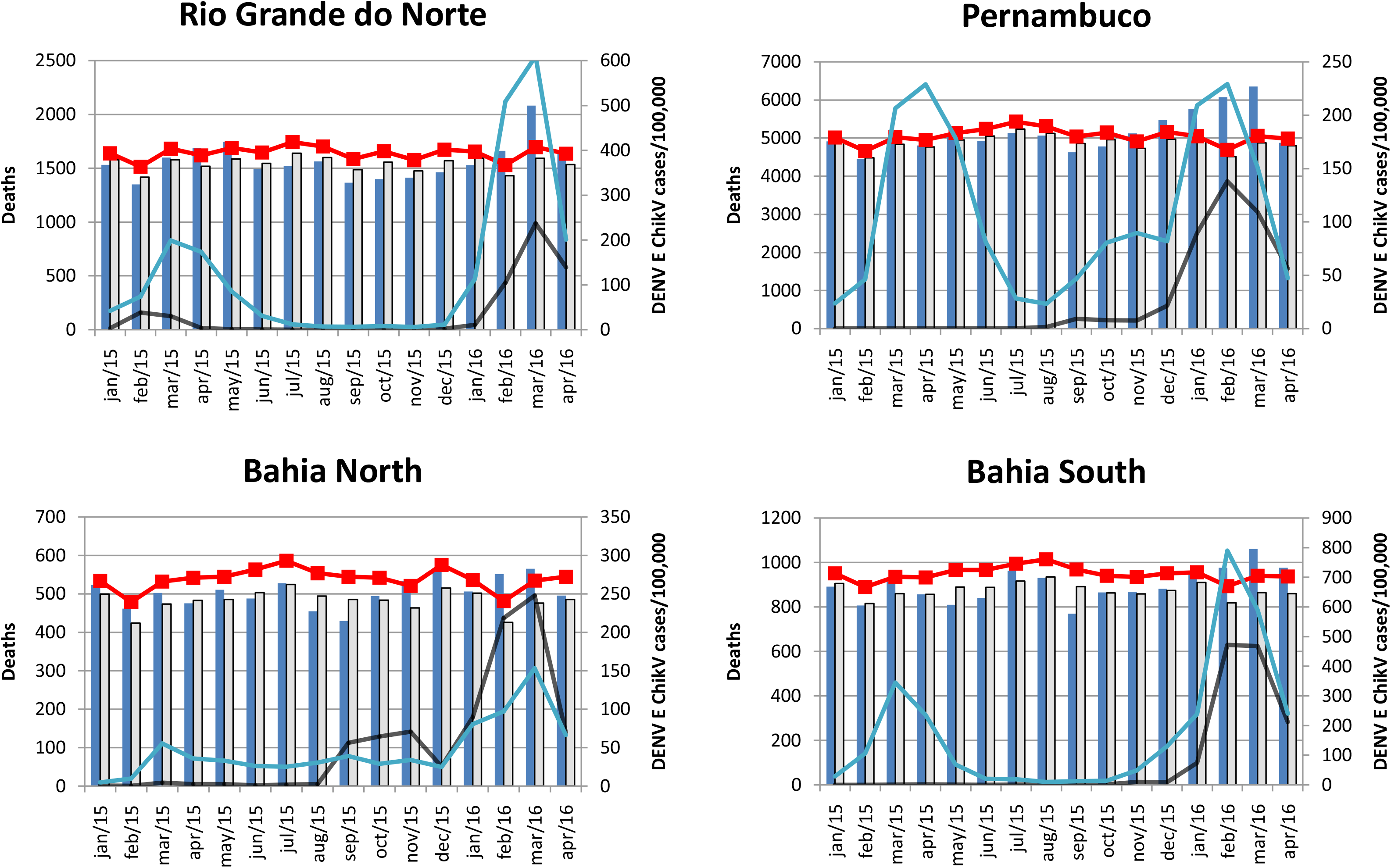
Change in mortality rate (deaths/100.000 inhabitants) by age group in Pernambuco, Rio Grande do Norte and Bahia (two regions), during chikungunya epidemics from 2015 to 2016. The asterisks indicate the age groups in which change occurred beyond the confidence interval (99%).

**Table 2.**
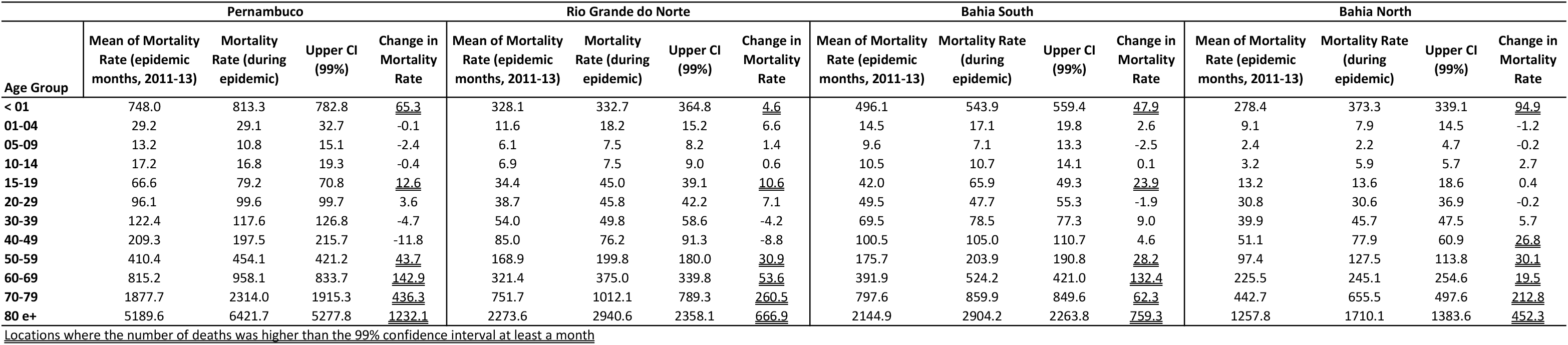
Observed, expected all-causes mortality rates (deaths/100.000 inhabitants) and excess mortality by age group in in Pernambuco, Rio Grande do Norte and Bahia (two most affected regions, North and South).

## Discussion

The understanding of the actual chikungunya burden depends on a better understanding of the mortality associated with this arbovirus. These findings show excess mortality during chikungunya epidemics in three most affected states in Brazil. The correlation between the incidence of chikungunya and increase in all-cause mortality reinforce the possibility that chikungunya virus has an impact on mortality rates. Similar results have already been found in other chikungunya epidemics in the past decades(2–4).

Since the chikungunya arrive in Americas, the virus has been identified in 44 countries and territories, leading to more than 2 million cases and 403 deaths(6). In Brazil, the first case was confirmed in September 2014; however, the worst outbreak occurred in 2016, with 236,287 reported cases and 120 officially confirmed deaths(5). The Brazilian Epidemiological Surveillance System detected far fewer deaths compared to our estimates in Pernambuco(54 deaths), Rio Grande do Norte(19 deaths), and all of Bahia (5 deaths)(5). Mortality identified by the passive surveillance system was approximately 50 times lower than estimated from the excess of deaths. These differences may be due to failure to diagnose and difficulty recognizing severe forms of chikungunya(1). Deaths caused by chikungunya occur more frequently in the elderly as a complication of preexisting diseases as reported in other hospital-based studies(7). The present study showed that the increase in mortality was higher than the 99% confidence interval in all regions in the age groups above 50 years. In fact the elderly are known to be the age group with the greatest risk of severe forms and deaths due to chikungunya(7). Because chikungunya is a disease not yet well known by health professionals in Brazil, the association of exacerbation of the underlying disease with chikungunya may have been underestimated leading to underreporting. In the age range between 15 and 19 years there was an increase in mortality rate above the 99% confidence interval in 3 of the 4 regions studied. There was also an increase greater than 99% confidence interval in the mortality rate in children under 1 year. These latter findings had not been previously described and need to be better understood however it is not possible to exclude the possibility that the excess mortality in these age groups was related to the circulation of zika virus.

The results of time-series studies must be carefully analyzed because of the method’s own difficulties, these difficulties include time-coincident confounding factors. The periods of excess mortality observed do not coincide exactly with the seasons of influenza in the states of the Northeast region. Influenza AHIN1 was the most frequent viral subtype in 2016 and has not been found to be associated with high elderly mortality(5,8). Zika virus was not a reportable disease in 2015 there is no official data to calculate the IR of zika virus in that year, but even so the mortality associated with zika is admittedly low (9). Other factors could be associated with excess mortality in that period; however, no changes in the epidemiological profile of other diseases or events in the regions studied were reported. Furthermore, the overall arbovirus incidence may be underestimated due to under-reporting by professionals. The impact of arbovirus co-circulation is still poorly understood. The interaction of arboviruses, could theoretically result in more intense viremia or other immunological alterations.

Considering the magnitude and rapid territorial expansion of chikungunya transmission, we believe that global health systems should be prepared to manage the severity of the disease and associated deaths, not only acute and chronic joint disorders that have been the main concern related to chikungunya historically. Excess mortality is a useful indicator to quantify the impact of a health-related event and has traditionally been used to describe the increase of deaths during influenza season and extreme climates events(8,10). All causes mortality during chikungunya epidemics could be monitored as a strategic tool, beyond individual case reporting to the epidemiological surveillance system, to assess excess mortality and the overall burden of chikungunya.

